# The Taurus Mountains, the hotspot of Western Palearctic biodiversity, is under danger: marble quarries affect wildlife

**DOI:** 10.1101/2023.11.02.564896

**Authors:** Tamer Albayrak, Tamer Yılmaz

## Abstract

The Taurus Mountains in the Mediterranean Coastal Basin, considered a biodiversity hotspot, have a rich biodiversity in the Western Palearctic. The number of marble quarries in the Taurus Mountains has dramatically expanded over the past ten years. The objectives of this study are to (i) determine the impacts of quarrying on wildlife; (ii) determine the potential impacts of quarrying on the future of Taurus. A total of 57547 photos and video images were analyzed on 5447 photo-trap days in two areas, the marble quarries and the control areas. Using 97 randomly selected marble quarries, the area they cover and their annual growth rates were determined. The most commonly seen animals were the wolf, fox, lynx, and wild boar in the control area, and the jackal and hare in the marble quarries (*p* < 0.001). Additionally, we found a significant positive correlation between the distance of the geographical center of the marble quarries and the number of dates of Wolf, Fox and Wild Boar and a negative significant correlation for hare (*p* < 0.05). A positive correlation was found between the area of marble quarries and the duration of operation (*R* = 0.89, *p* < 0.00). The waste from quarries, which makes up 79.7% of the total land used for this purpose, is the greatest cause of habitat degradation. According to calculations, even if no new marble quarries are built as of right now, 7.14% of the Taurus may have disappeared by the year 2027, and by the year 2032, 8.25% of the Taurus ecosystems may have disappeared completely. The Taurus Mountains, a center of Western Palearctic biodiversity, are being threatened by the marble quarries. This study advances our knowledge of how marble quarries may affect wildlife. New strategies must be developed as soon as possible to protect the Taurus Mountains, the hotspot of the Mediterranean basin.

**Summary Text for Online Table of Contents:** We conducted a one-year photo trap study on the effects of marble quarries on wildlife in the Taurus Mountains, a significant biodiversity hotspot. Our research showed that wild animals prefer areas without marble quarries, and their density rises as they get farther from the quarry’s core. Photograph by Tamer Yılmaz

## Introduction

The diversity of its plants and animals makes Anatolia one of the Western Palearctic’s (WP) high biodiversity zones. Anatolia was a key refuge for many WP species during the last ice age and includes a variety of habitats from sea level to 5000 meters above sea level. The animals and plants of the northern regions of WP, which are not part of Anatolia, moved to the Anatolian refuges during the Ice Age. They evolved there, some of them stayed in Anatolia, and some of them moved to Europe to recolonize the European fauna after the Ice Age (Hewitt 1999). Due to this kind of influences, Anatolian biodiversity has increased by 169 mammal (Özkurt and Bulut 2021), 491-512 bird (Kiziroğlu 2015), 139 reptile (Ilgaz 2019) and 35 amphibian (Kurnaz 2020), 33820 insect (Tezcan 2020), and 12000 plant (Şenkul and Kaya 2017) species. A large proportion of them live in the Taurus Mountains, which extend along the Mediterranean coast in WP (Ciplak 2003; Atalay 2006; Albayrak *et al*. 2012a). In addition, there are unique endemic species in the Taurus Mountains (Şenkul and Kaya 2017), Taurus-specific genetic structures of widespread populations, e.g., of mammals (Gür *et al*. 2018), birds (Albayrak *et al*. 2012b), amphibians (Göçmen and Akman 2012), relict species from the last glacial period e.g. Orthoptera, (Ciplak 2008). In addition, newly discovered plants (Yildirim and Tekşen 2021) and animals (Lohaj and Anlaş 2021) are found in Taurus.

Recently, a significant amount of biodiversity has been lost due to various reasons such as industrialization and rapid increase in human population (McDonald *et al*. 2020). In recent decades, some species such as the Falkland wolf (Sillero-Zubiri 2015) and Pinta Giant Tortoise (Cayot *et al*. 2016) have disappeared from the wild. According to IUCN, 26% of mammals, 14% of birds, 41% of amphibians are considered threatened species (IUCN 2021). Anthropological impacts in rainforests and Mediterranean basins, which are hotspots of world’s biodiversity, threaten many species, including potential species we have yet to discover. Habitat destruction and fragmentation, agricultural monocultures are causing natural habitat collapse and population declines. Marble quarries have the same effect, causing habitat destruction and fragmentation. Over time, this situation leads to the extinction of species. Therefore, monitoring wildlife is extremely important for biodiversity conservation, and the photo-trapping method is extremely useful for this purpose.

The photo-trapping method is useful for detecting wildlife, especially cryptic species such as Caracal, *Caracal caracal* (Giannatos *et al*. 2006), Wolf, *Canis lupus* (Albayrak 2011), and Monk Seal, *Monachus monachus* (Gucu *et al*. 2009). In addition to detecting the species with a systematic photo-trapping study, daily and seasonal activity times can also be determined (Naidenko *et al*. 2021). To identify wildlife in Anatolia, photo-trapping studies were conducted in Beydağları (Albayrak *et al*. 2012a), Yenice Forest (Can and Togan 2009), and deciduous forests (Çoğal and Sözen 2020).

Marble quarries cause habitat destruction by removing the soil layer and exposing the stone layer. It is not precisely known whether the number of marble quarries in Turkey has increased in the last decade, but exports amounted to $ 4.6 million in 2000 (about 500 quarries) and increased 370-fold to $ 1.7 billion in 2020 (about 1500 quarries) (Ticaret Bakanlığı 2021). Marbele quarries showed a very serious increase of 891.44% (from 148.41 hectares to 1577.52 hectares) between 1995 and 2020 in a local area of Taurus Mountain (Tercan and Dereli 2021). Marble quarries are operated with low productivity due to the rock structure of the Taurus. In addition to the irreversible destruction of the habitat where the marble quarries are operated, wildlife is expected to be affected by human activity and noise pollution caused by the 24/7 operation of the quarries. The Taurus Mountains, one of the most important areas of the Mediterranean region, which is considered a hotspot of Western Palearctic biodiversity, may be threatened by the rapid increase of marble quarries and wildlife can be affected by these activities. Unfortunately, to the best of the authors’ knowledge, there are no studies investigating the influence of marble quaries on wildlife. Therefore, the objectives of this study are to (i) determine the impacts of quarrying on wildlife; (ii) determine the potential impacts of quarrying on the future of Taurus. To achieve these objectives, we first determined the number of current marble quarries and then examined the impact of these quarries on wildlife. More than 50000 photo-trap images were analyzed in the area where the quarries are located and the adjacent control area to determine if the quarries are affecting wildlife.

## Method

### Study area and design

The study was conducted in the western Taurus Mountains (Taurus). To determine the effects of marble quarries on wildlife, two adjacent areas, each approximately 120 km^2^ and with the same habitat type and elevation, were selected as the marble quarry area (MQA) and control area (CA). MQA is located in the Dumlu Mountains, where 51 marble quarries are active, and CA is located in the Söğüt Mountains, where there are no marble quarries (Figure 1). The occupied areas of the randomly selected 97 active marble quarries were calculated using Google Earth, and Google Earth Timelaps was used to describe the year in which operations began in the western Taurus.

**Figure 1.**
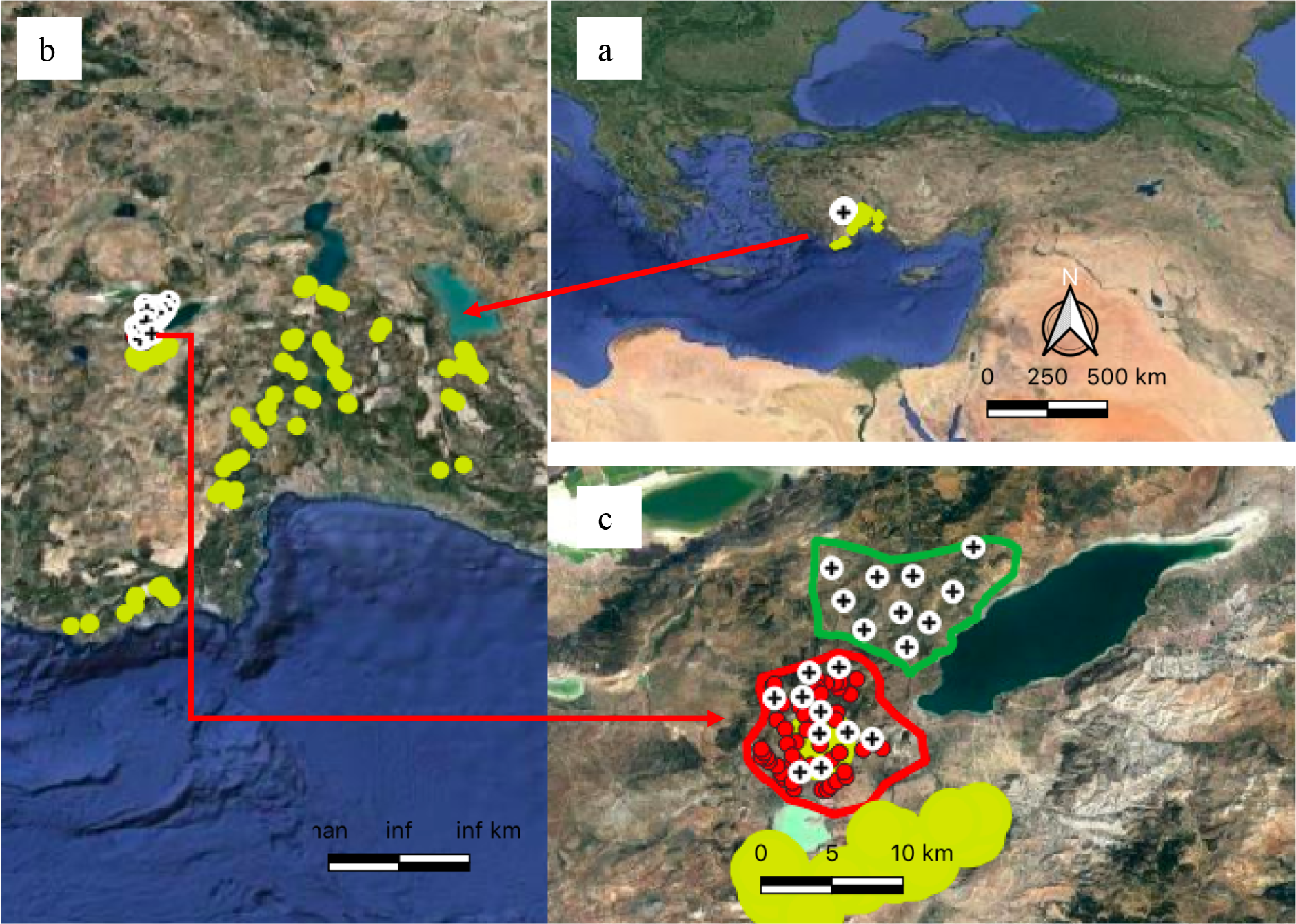
Western Taurus is the study area and the areas where the photo-trapping study will be conducted. a: Study area; b: yellow dots are randomly selected marble quarries in western Taurus; c: Areas where the photo-trap study is conducted, red dot are marble quarries in MQA and green line is the control area, plus signs are the photo-trap locations.

### Photo-trap and target species

To determine photo-trap locations, each study area was divided into 30 quadrats of 4 km^2^, and ten quadrats were selected to represent the entire study area on the map. A total of 20 Bushnell Trophycams and Reconyx UXR6 phototraps, ten each, were placed on animal track transit routes in MQA and CA. The photo traps, two photos and ten seconds of video per trigger, were operated on 399 days between 01/12/2015 and 02/01/2017. The result of the analysis of 57547 photo and video images from 5447 photo-trap days was 16,995 mammal data obtained by combining images taken simultaneously. Six target species were evaluated in the study, including Wolf (*Canis lupus*), Jackal (*Canis aureus*), Fox (*Vulpes vulpes*), Lynx (*Lynx lynx*), Wild Boar (*Sus scrofa*), and Hare (*Lepus europaeus*).

### Data analyses

The average annual growth rate of the marble quarry is calculated by dividing the area covered by the quarry by the year of its operation. The areas that will be covered after 5 and 10 years were calculated using the average annual growth rate (n = 97). To determine the differences in prevalence of target species between the marble quarries and the control areas, a t-test was performed. Central location was determined using the coordinates of the 51 quarries in the MQA. To understand the impact of the quarries on wildlife, a correlation analysis was conducted between the prevalence of target species and their distance from the central location.

A correlation analysis was also conducted to determine the relationship among the target species. The numbers reported in the analyses are the total number of data collected in a day, indicating the frequency of target species activity, not the total number of individuals in the field. All statistics were generated in R Studio (R Core Team 2021) and SPSS 17 software (Kalaycı 2010).

## Results

### Marble quarries

In general, there were few marble quarries in Taurus in 2005, but the number of quarries increased rapidly after 2008. A significant positive correlation was found between the total number of years and the managed area of the quarries (*R* = 0.89, *p* < 0.01; Figure 2), a positive correlation between the year in which the marble quarries were put into operation and the managed area (*R* = 0.18, *p* = 0.07; Figure 2). The first of the 97 selected marble quarries started operation in 2005, 91 of them were in operation in 2011, in 2016 all quarries were in operation and used a total of 1868 hectares, and a total area of 3164 hectares was calculated that would be used by 97 marble quarries in April 2021 (Figure 2). It was calculated that the average area of a marble quarry from the year of activity of 97 selected marble quarries from western Taurus in the 5th, 10th, and 15th years is 13.34 ± 15.89 (n = 97), 24.88 ± 29.66 (n = 91), and 49.20 (n = 1) hectares, respectively. If this increase continues at the calculated rate of increase, it is calculated that the 97 marble quarries selected as a sample will destroy the habitat of an area of 8541 hectares in 2027 after five years and 19540 hectares 10 years later in 2032, at a rate of increase of 18% per year (Figure 2). It was found that only 20.3 ± 6.6 % of the total area where the quarries caused habitat destruction was the area where the marble blocks were quarried, while 79.7 ± 6.6 % was the area where the waste produced during the quarrying of these blocks was disposed of.

**Figure 2.**
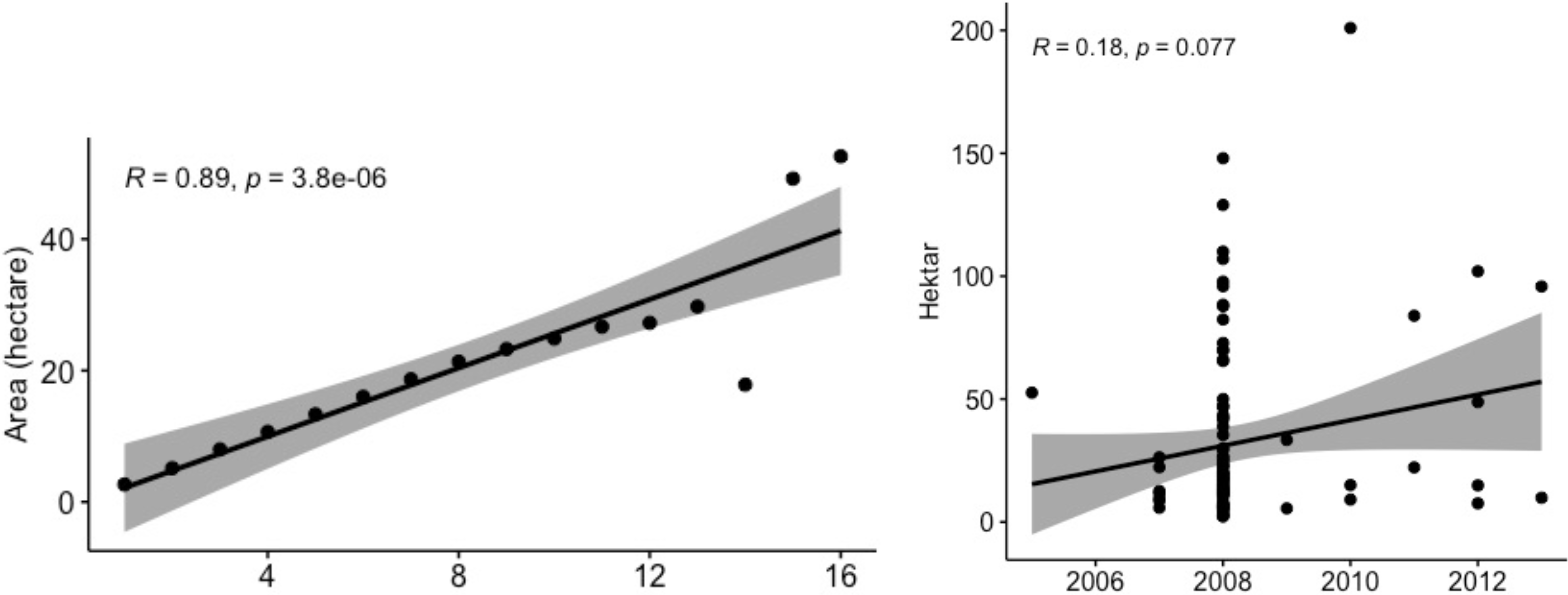
Growth rate of marble quarries (n=97); correlation analysis between years of operation and covered area (left). Correlation analysis between the uptake of the activity and the current cultivated area of the quarries (right).

### Photo traps

To determine the impact of marble quarries on wildlife, we analysed a total of 57547 photo and video images from a total of 5447 photo-trap days in the MQA and CA. We identified 12 mammal species from 16995 data. Of these data, 13441 included the target species Wolf (*Canis lupus*), Jackal (*Canis aureus*), Fox (*Vulpes vulpes*), Lynx (*Lynx lynx*), Wild Boar (*Sus scrofa*), and Hare (*Lepus europaeus*). In addition to the target species, Beech Marten (*Martes foina*), Pygmy Weasel (*Mustela nivalis*), European Badger (*Meles meles*), Southern White-breasted Hedgehog (*Erinaceus concolor*), Anatolian Tree Squirrel (*Sciurus anomalus*) and Williams’ Jerboa (*Allactaga williamsi*), which was recorded for the first time for Burdur province, were also recorded.

Looking at the monthly maximum numbers of the target species in MQA and CA, we find that their abundance varies depending on the month. During the summer months, the abundance of target species generally decreased, with the exception of feral hog. It can be seen that seasonal densities are similar between Wolf and Wild Boar and between Lynx and Hare (Figure 3).

**Figure 3.**
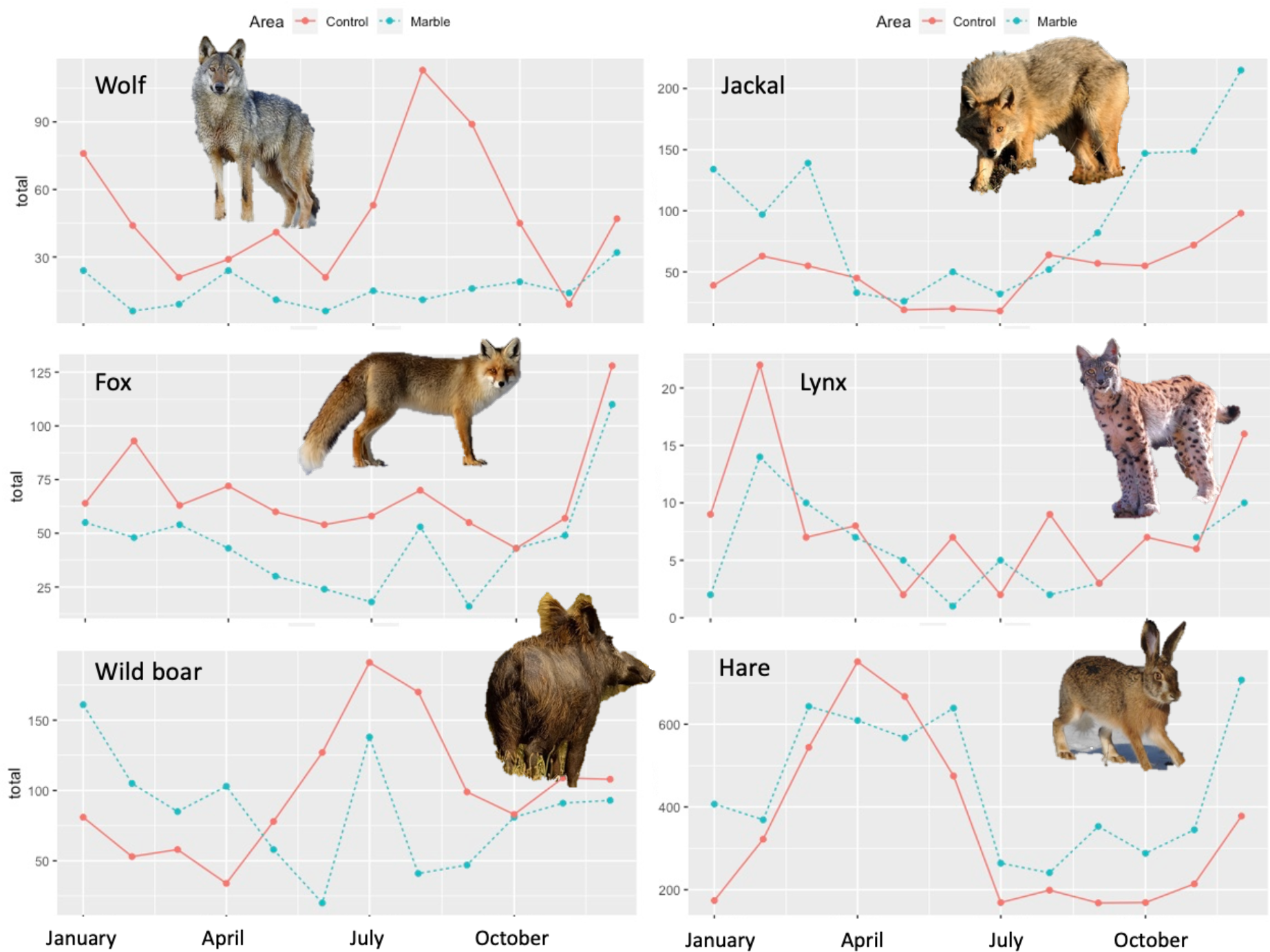
Total maximum numbers of monthly target species associated with areas.

When examining the daily activity times of the target species in relation to the areas, it was found that the species were generally active at night and had low activity between 10:00 and 15:00 (Figure 4). Wolf, Fox, and Jackal appeared to be more active during the day in the control area, but no statistical difference was found between the two areas (*p* > 0.05; Figure 6).

**Figure 4.**
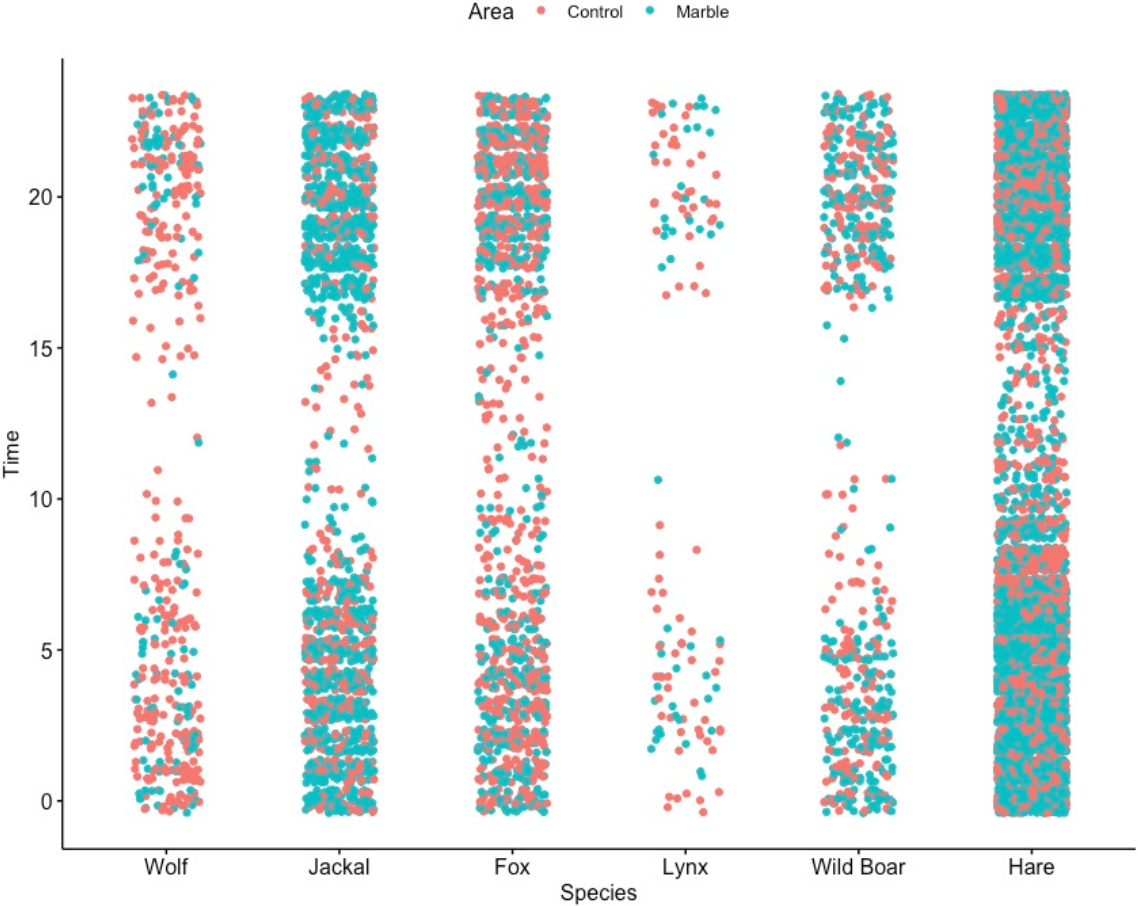
Daily activity times of target species, depending on areas.

Four predators and two prey species compared by total number of individuals recorded in a day are listed in Table 1 and Figure 3. Wolf, Fox, Lynx, and Wild Boar were more abundant in control areas; these differences were statistically significant for Wolf and Fox (*p* < 0.001). Hare and Jackal were statistically more common in the marble quarry area (*p* < 0.001; Table 1). A positive correlation was found between some target species, while a negative correlation was found between wolf and hare when two areas were evaluated together (Figure 5).

**Table 1.**
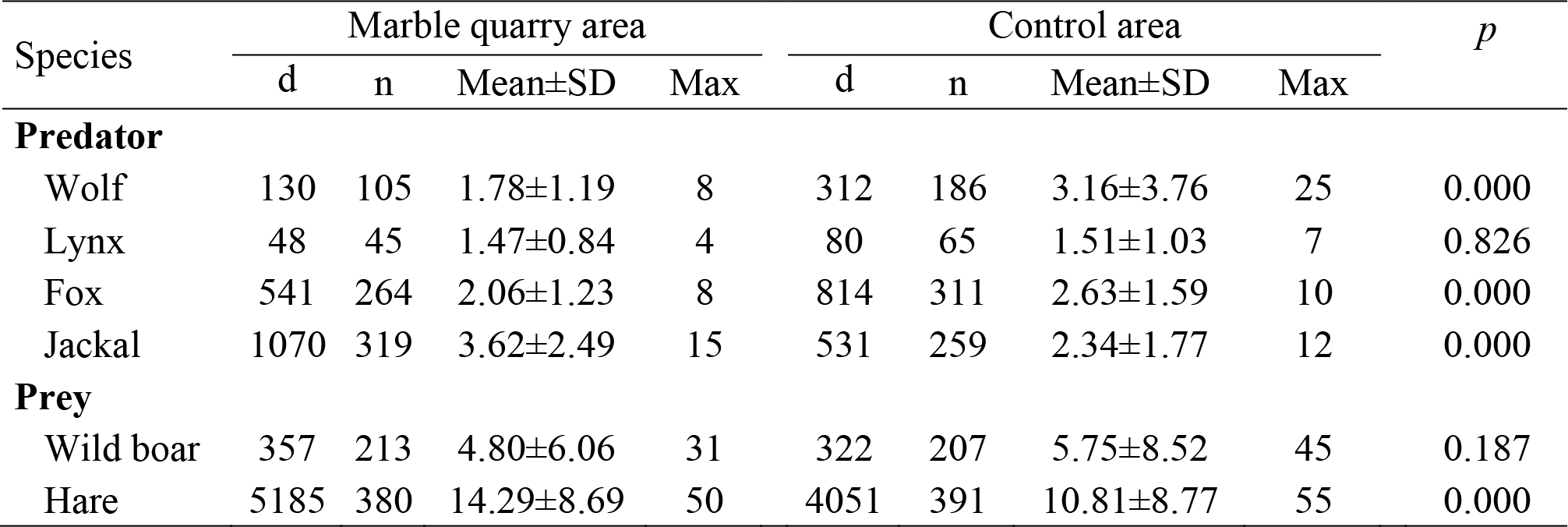
Statistical comparison of total number of individuals of target species recorded in a day. d: number of data; n: Number of days with data.

**Figure 5.**
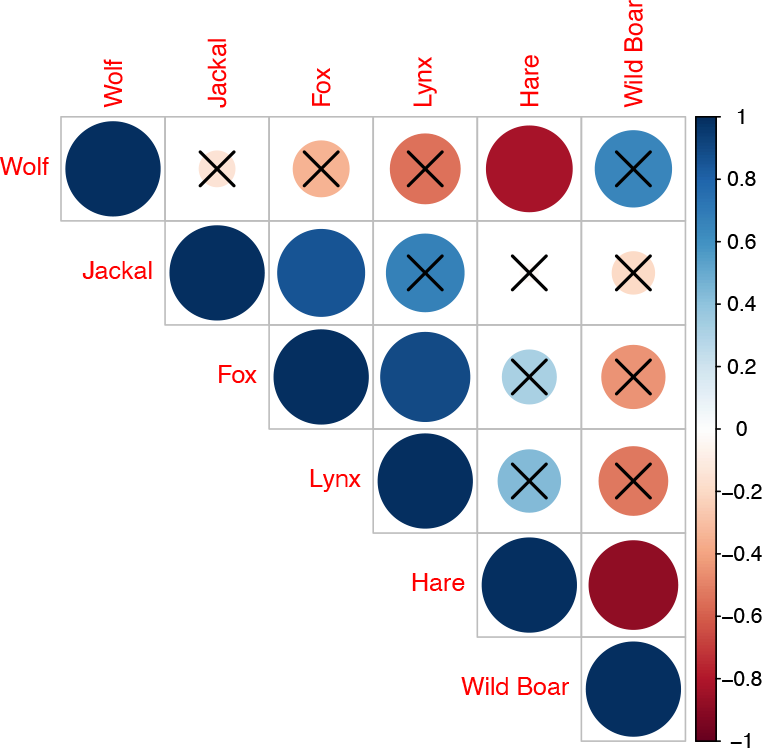
Pearson correlation between target species. The colour indicates the *R* value, the size of the circle the *p*-value and the cross indicate nonsignificant (*p* > 0.05).

We found a positive correlation between the distance of the photo-trap site from the centre of the quarries and the occurrence of Wolf, Fox, and Wild Boar. The Wolf (*R* = 0.21, *p* < 0.01), Fox (*R* = 0.09, *p* < 0.01) and Wild Boar (*R* = 0.09, *p* < 0.05) were affected by the marble quarries and were found at a greater distance from the quarry centre (Figure 6). While there was no statistical correlation for Jackal and Lynx, a negative correlation was found for hare (Figure 6).

**Figure 6.**
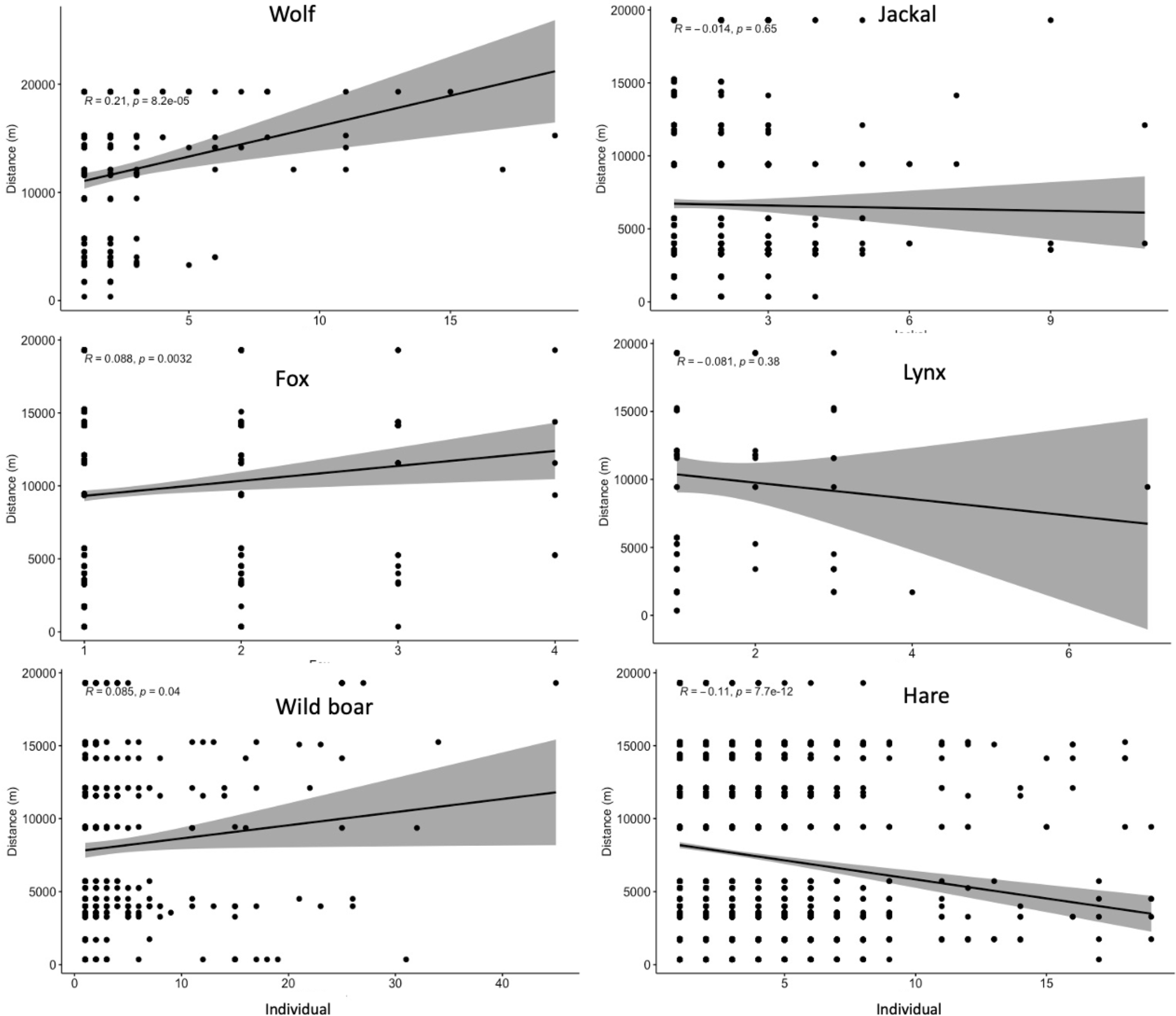
Correlation between distance of target species to quarry centre and daily total number of individuals.

## Discussion

### (i) habitat destruction by marble quarries

Marble activities have attracted the attention of people in Turkey at all times, from the Roman Empire period until today (Long 2017). Since the rock structure of the Taurus Mountains (Taurus) consists of limestone suitable for marble quarrying, marble quarries, which are open-pit mines, have become the focus of activities (Ticaret Bakanlığı 2021). In order to mine the marble blocks, the surface layers, soil and vegetation, must be removed and the unbroken rocks on the ground must be cut. Due to the fragmented rock structure of the Taurus Mountains, marble quarries operate at low productivity and produce too much waste. More habitat is destroyed due to the way of waste disposal. It was found that only one of five (20.3 ± 6.6 %) of the total area was the area where the marble blocks were quarried, while four of five (79.7 ± 6.6 %) was the waste produced during the quarrying of these blocks. The operation of marble quarries produces too much waste, which is indiscriminately disposed of from the slopes due to the low productivity of the quarries. Habitat destruction in quarries increases significantly with years of operation in western Taurus (Figure 2; *R* = 0.89 *p* < 0.01). In addition, a positive relationship was found between the area covered by quarries and the year of construction. Although it is not statistically significant (Figure 2; *R* = 0.18 *p* = 0.077), the newly opened marble quarries occupied more area than the old quarries.

### (ii) Impact of marble quarries on wildlife

Species have been found to be negatively affected by the alteration of an area’s natural habitat by changing species diversity (Zhu *et al*. 2004; Dumbrell *et al*. 2008) and the populations of mammals (Schmiegelow and Mönkkönen 2002; Bowler *et al*. 2019), birds (Schmiegelow and Mönkkönen 2002; Bowler *et al*. 2019), reptiles (Stanford *et al*. 2020; Cabrera 2021), and amphibians (Schmiegelow and Mönkkönen 2002; Bowler *et al*. 2019). The community changes when habitat is altered, but nearly all species disappear in the area when habitat is destroyed. Marble quarries destroy habitat and lead to habitat fragmentation. In this context, we found that the area covered by marble quarries is equivalent to the habitat fragmentation caused by quarries, but much larger.

We analysed 57547 photos and video images on 5447 photo-trap days, for a full year in the quarry area and in the control area without quarries, which has similar habitat and elevation, to determine the impact of the marble quarries on wildlife. Although the target species Wolf, Jackal, Fox, Lynx, Wild Boar, and Hare were detected in both the marble quarries and the control area, Wolf, Fox (statistically significant *p* < 0.05), Lynx, and Wild Boar were more abundant in the control area (Table 1). In addition, the occurrence of Wolf, Fox, and Wild Boar increased statistically the farther they were from the geographic centre of the marble quarries (*p* < 0.05; Figure 6). Habitat destruction and fragmentation caused by marble quarries harms wildlife and human activities such as noise, suggesting that species move as far away from quarries as possible.

Jackal and Hare were found to be more abundant in the marble quarries than in the control area. It is thought that this is due to the fact that jackal usually feeds on human waste caused by human activities. Ćirović et al. (2016) found that 71.8% of the Jackal’s diet was human waste. The fact that the jackal was more abundant in the Marble Quarries area is probably due to the fact that its predators, Wolf, Fox and Lynx, are less abundant in this area compared to the control area, as well as to the absence of illegal hunters, since the Marble Quarries are in operation around the clock.

### (iii) Potential impact of marble quarries on the wider Taurus, the hotspot of Mediterranean biodiversity

The Mediterranean Coastal Basin is a region of high biodiversity in the Western Palearctic (Myers *et al*. 2000); within this region, the Taurus Mountains form the Western Palearctic hotspot. The main reasons why the Taurus Mountains are a biodiversity hotspot are a mountain range that rises up to 3000 m above sea level, its unique microclimate, and its habitats that range from scrub to alpine zones. Moreover, the southern part of the Taurus served as a vital refuge for many animal species during the last ice age. They evolved and after the ice age, the species expanded their range from this refuge and restored the European biogeography. The Taurus Mountains harbor not only endemic species (Göçmen and Akman 2012) under the influence of the Ice Age, but also specific genetic structures of some widespread species such as Fallow deer (Baker *et al*. 2017), and Kruper’s Nuthatch (Albayrak *et al*. 2012b). The presence of newly identified plant and animal species is an important indication that the biodiversity of Taurus is not yet fully known. This could be due to the deep valley and difficult to access cliffs of Taurus. The existence of orchid species living only in a single valley of Taurus (Hürkan *et al*. 2015; Deniz *et al*. 2018), seven species and 21 subspecies of the genus *Lyciasalamandra* from southern Taurus, (Göçmen *et al*. 2013; Mezzasalma *et al*. 2021) and a specific clade of populations due to refugium effect on population (Kornilios *et al*. 2011; Albayrak *et al*. 2012b) are important examples of how specific species and genetic structure have evolved in Taurus.

Taurus’ unique limestone habitats are being destroyed by rapidly increasing marble quarries in the last decade. Our study found that target mammal species are being impacted by direct marble quarries and are moving as far away as possible from the geographic center of the quarries. While the lower and southern parts of the Taurus are already under severe pressure from tourism and human settlement, the northern and higher parts are also under severe pressure from marble quarries. The main reason for habitat destruction is the indiscriminate burial of waste from the quarries. Tercan and Dereli (2021) found that forest and semi-natural areas are generally allocated to marble quarries and the quarries, which is 4.69% of total area, showed a serious increase of 891.44% (average 164.48% in five years) between 1995 and 2020 in Burdur province. According to our calculations using growing rate of Tercan and Dereli (2021) (*R* = 0.98 *p* < 0.01 between percentage of MQA in total area and year), it is expected that even if no new marble quarry is established as of today, because the marble quarries have been established throughout the Taurus Mountains, 7.14 % of the Taurus in 2027 and ten years later, in 2032, the 8.25 % of the Taurus habitats may have disappeared. This situation will lead to both the extinction of locally endemic species and the extinction of the genetic structures of many species specific to the Taurus Mountains.

### (iv) Recommendations

When issuing operating permits for the marble quarries, new methods should be created. We have found that the main problem of habitat destruction is the waste from the quarries, which accounts for 79.7% of the total area used for this purpose. To protect the Taurus, the hotspot of Mediterranean biodiversity, marble quarries with low productivity should not be allowed, and quarry waste should not be allowed to be dumped indiscriminately on the slopes. The waste should be collected in a designated area. When issuing and controlling licenses for quarries, scientific studies should be conducted that take into account the biodiversity of the region. New strategies must be developed as soon as possible to protect the Taurus Mountains, the hotspot of the Mediterranean basin.

### Conclusion

Habitat destruction and fragmentation of marble quarries were found to negatively impact wildlife. Our hypothesis was confirmed by the target species, with the exception of the jackal, which feeds mainly on human waste. Marble quarries, which have multiplied in the last decade, are causing habitat destruction and fragmentation in the Taurus Mountains. Marble quarries, which multiply like cancer cells that have metastasized in the last decade, pose a significant threat to the Taurus Mountains, the hotspot of Western Palearctic biodiversity.

## Acknowledgments

The experiments comply with the current laws of the country in which they were performed. We thanks to Yusuf Çınar who drove the marble quarries areas in google earth.

## Author contributions

TY conducted fieldwork, edited data, assisted in article writing, TA analyzed data, wrote article.

## Data availability

The raw data are available from the corresponding author upon reasonable request.

## Declarations

### Ethical approval

Not applicable.

### Consent to participate

This study not involve human subjects therefore consent to participate was not required

### Consent to publish

The authors give Biodiversity and conservation authority to publish this manuscript.

